# Optogenetic Silencing of Medial Septal GABAergic Neurons Disrupts Grid Cell Spatial and Temporal Coding in the Medial Entorhinal Cortex

**DOI:** 10.1101/2023.11.08.566228

**Authors:** Jennifer C. Robinson, Johnson Ying, Michael E. Hasselmo, Mark P. Brandon

## Abstract

The hippocampus and medial entorhinal cortex (MEC) form a cognitive map that facilitates spatial navigation. As part of this map, MEC grid cells fire in a repeating hexagonal pattern across an environment. This grid pattern relies on inputs from the medial septum (MS). The MS, and specifically its GABAergic neurons, are essential for theta rhythm oscillations in the entorhinal-hippocampal network, however, it is unknown if this subpopulation is also essential for grid cell function. To investigate this, we used optogenetics to inhibit MS-GABAergic neurons during grid cell recordings. We found that MS-GABAergic inhibition disrupted grid cell spatial periodicity both during optogenetic inhibition and during short 30-second recovery periods. Longer recovery periods of 60 seconds between the optogenetic inhibition periods allowed for the recovery of grid cell spatial firing. Grid cell temporal coding was also disrupted, as observed by a significant attenuation of theta phase precession. Together, these results demonstrate that MS-GABAergic neurons are critical for grid cell spatial and temporal coding in the MEC.

## Main Text

The medial entorhinal cortex (MEC) contains a diversity of spatially modulated neurons, that include grid cells which fire at regular spatial intervals to form a hexagonal pattern of spatial fields across an environment^1-3^. Current theories predict that these repetitive patterns support navigation and memory encoding by providing self-motion-based information represented in a metric of space^1,4-6^. The networks that support grid cell spatial periodicity include the anterior thalamic nuclei (ATN)^7^, the hippocampus^8^, and the medial septum (MS)^9,10^. The MS is critical for theta rhythm generation in the hippocampus and the MEC. Lesions or inactivation of the MS disrupt theta oscillations in the hippocampal and entorhinal networks^11-13^ and cause significant spatial memory deficits^14,15^. Importantly, the MS consists of three distinct cell types: GABAergic, glutamatergic, and cholinergic neurons^16^. It remains unclear which cell type supports grid cell spatial firing. Previous data suggest that MS-GABAergic neurons are the central pacemaker of theta rhythm activity (6-12 Hz) in the MEC and hippocampus, as demonstrated by their burst firing pattern at theta frequencies in-vivo^17^, their temporal lead ahead of hippocampal theta activity^18^, and their primary contact with hippocampal and entorhinal interneurons^19,20^ that pace theta oscillations^21^. Optogenetic activation of the MS-GABAergic subpopulation of parvalbumin (PV) neurons entrains hippocampal and MEC oscillations^22-24^, while silencing septal GABAergic neurons results in a large decrease in the power of endogenous theta oscillations during open field exploration and rapid-eye movement (REM) sleep^25,26^. It is well established that MEC grid cell periodicity relies on inputs from the MS^9,10^, but recent studies call into question the necessity of theta oscillations for grid cell periodicity^24,27^. Optogenetic activation of GABAergic subpopulation PV neurons in the MS paces ongoing oscillation well above theta range but does not disrupt the grid cell spatial pattern^24,27^. To more directly test whether MS-GABAergic driven theta oscillations support grid cell firing, we used optogenetic inhibition to selectively silence MS-GABAergic neurons to reduce endogenous theta oscillations to examine their contribution to both spatial and temporal coding of grid cells in the MEC.

### Selective inhibition of MS-GABAergic neurons in the MS decreases the power of theta oscillations and grid cell periodicity

To target MS-GABAergic neurons in the MS, Archaerhodopsin-3 variant ArchT^28^ was selectively expressed in the MS (Fig. 1a). Selectivity was driven by Cre-dependent adeno-associated virus injected into the MS of VGAT^cre^ mice (Supplementary Methods). An optic fiber was positioned above the MS for light delivery while both single-unit and local field potential (LFP) recordings were recorded in the MEC (Fig. 1b). Consistent with prior work^25,26^, MS-GABAergic neurons inhibition resulted in a clear decrease in the power of endogenous MEC theta oscillations (Fig. 1c). In contrast, Green Florescent Protein (GFP) control animals show a slight but non-significant increase in theta power during laser-on periods (Fig. 1d). The optogenetic inhibition protocol consisted of 30 second laser-on periods alternating with 30 second interstimulus intervals (ISI) repeated throughout the full recording session (Supplementary Methods).

**Figure 1.**
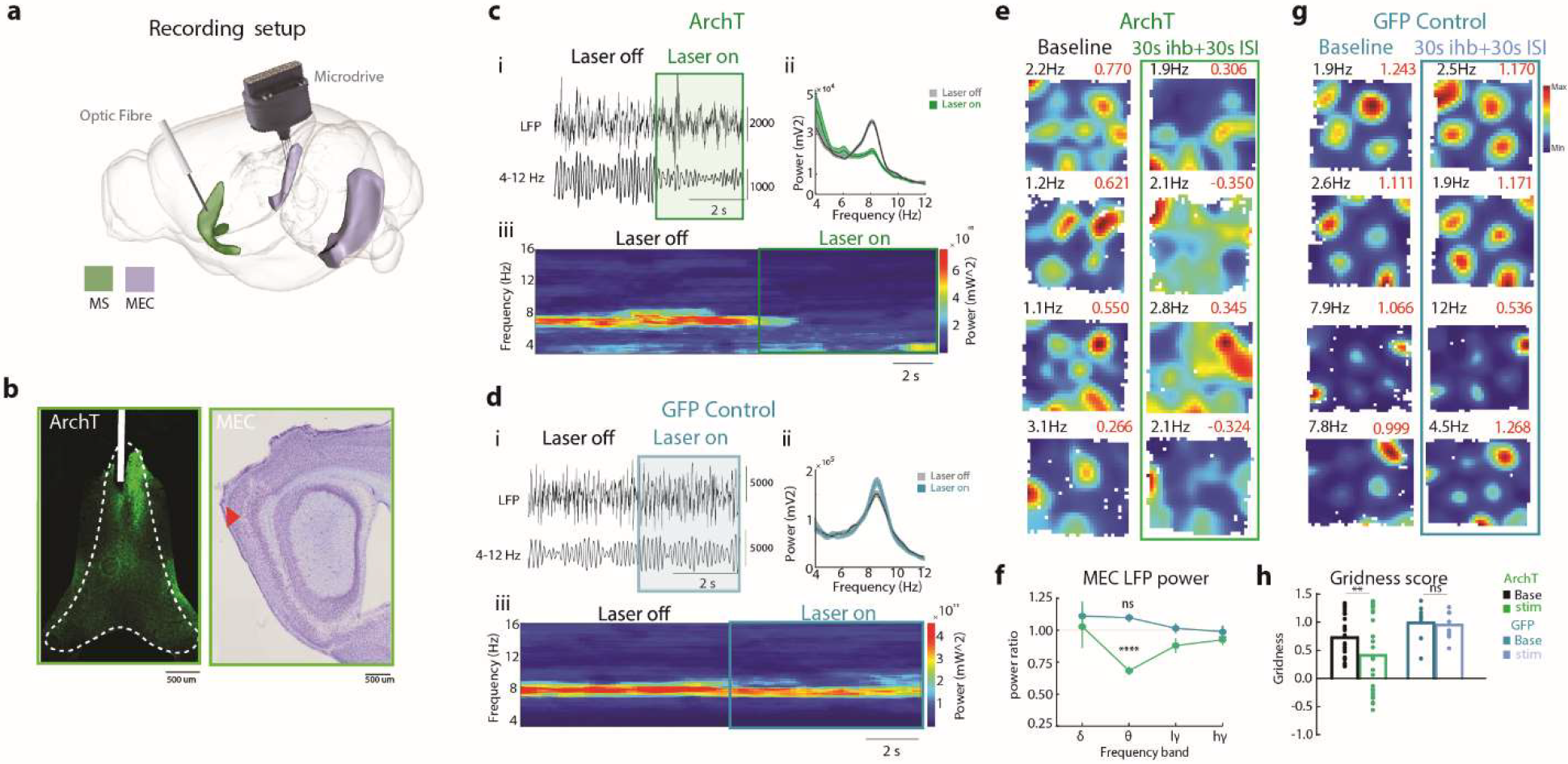
Optogenetic targeting of MS-GABAergic neurons with archaerhodopsin reduces endogenous theta oscillations and disrupts grid cell spatial firing. (a) Illustration of stimulation and recording set up with an optic fibre placed above the MS at a 5-6° angle and a 4 tetrode microdrive placed above the medial entorhinal cortex, illustration modified from https://scalablebrainatlas.incf.org/. (b, left) Example image of Virus expression of AAV-ArchT-GFP shown in green in the MS. (b, right) Example electrode track in the MEC. (ci) Example of raw (above) and filtered LFP (below) during laser-off and laser-on periods in animals injected with AAVdj-ArchT. (cii) Example power spectrum for full recording session with repeated 30s laser-on (green) and 30s laser-off periods (grey). (ciii) Example spectrogram corresponding to ci within theta range. (di) Example of raw (above) and filtered LFP (below) during laser off and laser on periods in animals injected with AAV2-GFP. (dii) Example power spectrum for full recording session with repeated 30s laser-on (blue) and 30s laser-off periods (grey). (diii) Example spectrogram corresponding to ci within theta range. (e) Example rate maps for cells recorded across baseline (left) and combined 30s silencing followed by 30s interstimulus interval (ISI) (right) with peak firing rate included in black and gridness score included in red above each rate map for ArchT group. (f) Proportion of 30s inhibition of delta, theta, low and high gamma power compared to 30s-ISI periods for ArchT (green) and GFP (blue). (g) Example rate maps for cells recording across baseline (left) and combined 30s silencing followed by 30s ISI (right) for GFP controls. (h) Gridness score for baseline and stimulation conditions for both ArchT and GFP groups. For all panels, ns = non-significant, \**P*<0.05, \*\**P*<0.01, \*\*\*\**P*<0.0001.

To quantify the extent to which grid cells were affected by MS-GABAergic inhibition, the gridness score (a measure of hexagonal spatial periodicity in the rate map; Supplementary Methods) was assessed during the baseline recording session, inhibition periods and ISI periods. The gridness score was significantly decreased during the stimulation recordings compared to baseline, [mean±SEM, Wilcoxon matched-pairs signed rank test; baseline mean 0.74±0.077, stim mean 0.43±0.14 P=0.0034, n = 25] (Fig. 1e and h, Supplementary Fig. S1), while no significant difference was found in GFP controls [means±SEMs, Wilcoxon matched-pairs signed rank test; baseline mean 1.01±0.10, stim mean 0.97±0.083, p= 0.94, n = 9] (Fig. 1g and h, Supplementary Fig. S2). Although the mean gridness score was significantly decreased, grid cells expressed a heterogeneity in their response: some cells maintained their periodic firing while others were more dramatically disrupted (see point distribution in Fig. 1h). It is possible that cells which maintain their spatial firing would require extended periods of MS-GABAergic inhibition to elicit a disruption in spatial periodicity. Alternatively, this subset of grid cells may be less reliant on theta oscillations, which is consistent with reports where grid cells maintained their periodicity firing without endogenous theta^24,29^. Considering that muscimol mediated septal inactivation results in both the loss of theta oscillations and periodic firing in all grid cells^9,10^, we predict that certain grid cells may require a more prolonged disruption of theta inhibition than was performed in this study. Consistent with previous findings^9^, head direction and other non-grid spatially modulated neurons were unaffected by the stimulations (Supplementary Figs. S3-S4). These results demonstrate that MS-GABAergic neurons facilitate grid cell spatial periodicity and provide supporting evidence for the essential role of theta oscillations in maintaining the spatial firing pattern of grid cells.

### Grid cell spatial firing does not recover immediately with the return of theta oscillations

To further examine the relationship between grid cell periodicity and theta oscillations, we split and concatenated recording sessions separately into inhibition and ISI periods (Fig. 2a). Compared to the baseline recording, grid cell periodicity was significantly disrupted during both the inhibition and ISI periods [mean±SEM: baseline mean 0.74±0.076, 30s inhibition mean 0.30±0.10, 30s ISI mean 0.31±0.12, Friedman test *P*<0.0001 with Dunn’s multiple comparisons test Baseline vs 30s-inhibition *P*<0.001, baseline vs 30s-ISI *P*<0.0001] (Fig. 2b). The absence of spatial periodicity in the 30s-ISI period when endogenous theta rhythms returned provides insights into the time that the grid cell network requires to recover and return to a normal grid firing pattern following MS-GABAergic inhibition. These data demonstrate that short bouts of theta oscillations are not sufficient to support grid cell spatial firing and network-wide dynamics may require more time to recover following MS-GABAergic inhibition.

**Figure 2.**
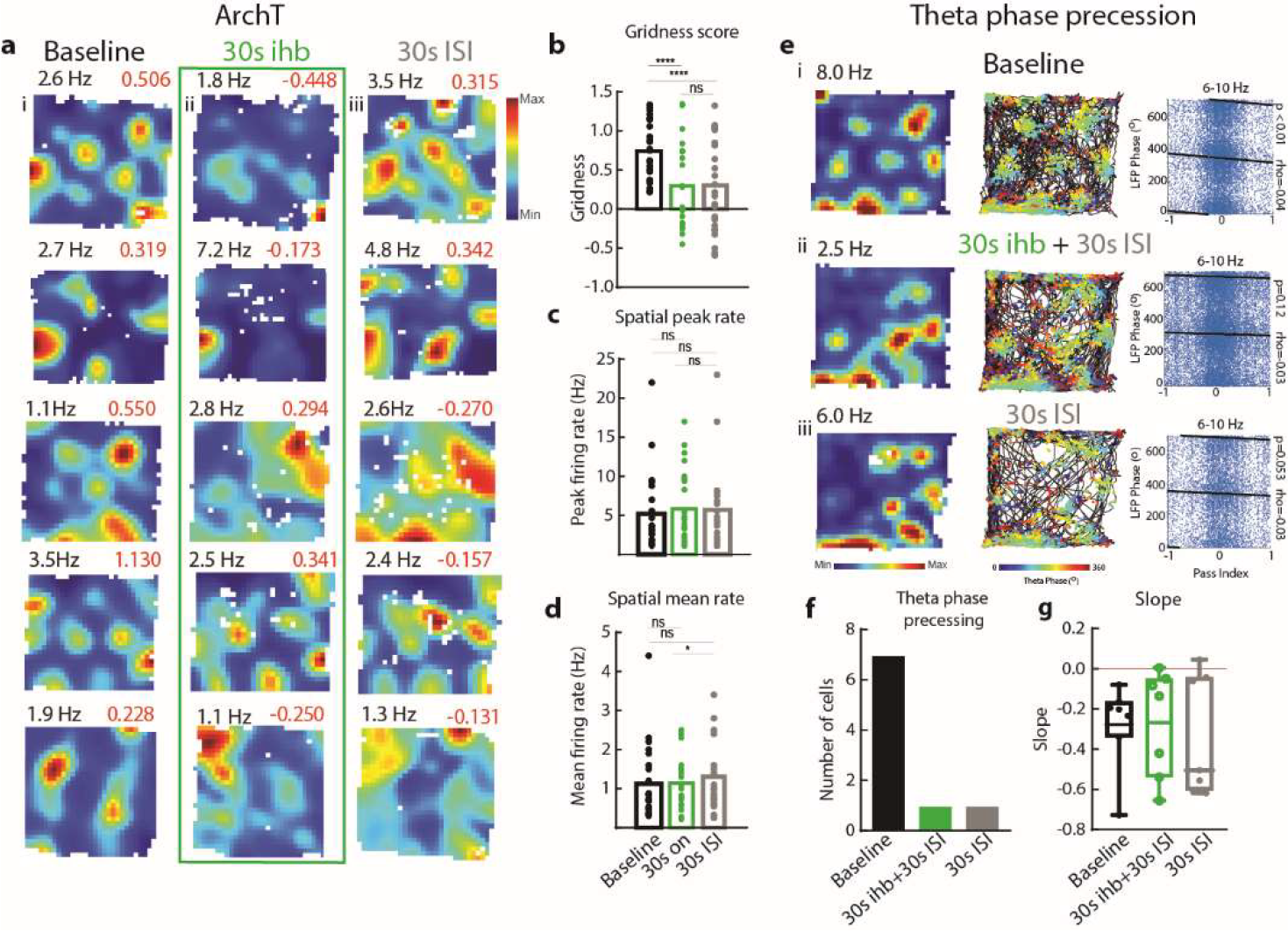
Optogenetic MS GABAergic inhibition disrupts grid cell spatial periodicity and phase precession during both 30s-inhibition and 30s ISI. (a) Example rate maps of grid cells in (i) baseline and separated into (ii) 30-inhibition (inhib) (iii) 30-ISI conditions. (b) Gridness score across baseline, 30s inhibition periods and 30s ISI periods. (c) peak firing rate (Hz) across baseline, 30s inhibition periods and 30s ISI periods. (d) mean firing rate (Hz) across baseline, 30s inhibition periods and 30s ISI periods. (e) Analysis of 2D precession data including rate map (left), trajectory plot (middle) and phase of spiking. Pass index values of -1 and +1 represent the entry and exit of a firing field. Regression line on the right indicates that the spike phase predicts the position of the animal within the field. P value of phase precession is shown to the right. Example unit ceased phase precessing during stimulation, shown by the loss of significant phase precession (*P* < 0.01). (f) Number of cells phase precessing during baseline and 30s laser-on 30s laser-off stimulations. We observed one neuron which continued to phase precess during the stimulation (*P* < 0.01). (g) Slopes for baseline, 30-inhib+30-ISI and 30s-ISI only periods. For all panels ns = non-significant, \**P*<0.05, \*\*\*\**P*<0.0001.

We further examined the general firing properties of grid cells and found no significant difference in peak firing compared to baseline [mean±SEM: Max firing (Hz) baseline mean 5.2± 0.97, 30s-inhibition mean 5.86±0.90, 30s-ISI mean 5.72±0.99, Friedman test P=0.61 with Dunn’s multiple comparisons test Baseline vs 30s-inhibition *P*=0.79, baseline vs 30s-ISI *P*>0.99] (Fig. 2c). However, mean firing was significantly higher during the 30s-ISI period compared to 30s-inhibition [mean±SEM: Mean firing (Hz) baseline mean 1.13±0.19, 30s-inhibition mean 1.14±0.13, 30s-ISI mean 1.31±0.16, Friedman test *P*=0.0164 with Dunn’s multiple comparisons test Baseline vs 30s-inhibition n.s., baseline vs 30s-ISI *P*=0.04] (Fig. 2d). This could be due to rebound activity following the inhibition period. Controls expressing GFP alone showed no significant changes in gridness score or firing rates compared to baseline (Supplemental Fig. S2).

### Modulation of MS-GABAergic activity disrupts grid cell temporal coding

As MS-GABAergic neurons contribute to ongoing theta oscillations in the MEC, periodic disruptions to rhythmic theta inputs to entorhinal interneurons may alter grid cell temporal coding in the form of impaired theta phase precession. Oscillatory interference and continuous attractor network models suggest that theta phase precession could allow grid cells to integrate spatial displacement based on self-motion while planning future paths as part of a path integration system^30-32^. Previous literature has shown that driving endogenous theta above theta range results in a disruption in grid cell theta phase precession^24^. To examine if theta phase precession is affected in grid cells during periodic optogenetic inhibition, we computed the strength of correlation between the distance travelled across a grid field and the theta phase of spiking (Fig. 2e). We observed that, of seven grid cells that exhibited phase precession during baseline recordings, six no longer showed phase precession when theta power is periodically disrupted during the 30s-inhibition with 30s-ISI recording sessions. We further examined if phase precession returned during the 30s-ISI periods and observed the same finding, that 6/7 grid cells exhibited no phase precession even with the return of theta oscillations (Fig. 2f). Interestingly, while the correlation between the spiking phase and distance travelled across a firing field phase precession was significantly disrupted, phase precession slopes remained negative in all except one neuron (Fig. 2g). Our results suggest that intermittent silencing of MS-GABAergic cells can disrupt the integration of self-motion-based cues via theta phase precession.

Given that grid spatial firing was disrupted during the 30s-inhibition period as well as the 30s-ISI period, we examined whether extending the duration of the ISI period would allow for recovery of grid cell spatial firing. In these experiments, we tracked neurons across four recordings: Baseline, 30s-inhibition with 30s-ISI protocol, ∼24 hrs recovery, and 30s-inhibition with 60s-ISI periods. Tracking the same cells with different interstimulus lengths revealed that a longer ISI period of 60s between inhibition periods allowed for the recovery of the grid cell spatial firing pattern (Fig. 3a and 3b, n=8, [mean±SEM: baseline mean 0.52±0.12, 30s Inhib+30s ISI 0.0036±0.14, 24 hrs post mean 0.75±0.13, 30s Inhib+60s ISI=0.33 ±0.20, Friedman test *P*<0.01 with Dunn’s multiple comparisons test Baseline vs 30s Ihb+30s ISI *P*<0.05, baseline vs recovery *P*>0.9999, baseline vs 30s Ihb+60s ISI *P*=0.2993]). Next, we examined the time-course of grid cell disruption and recovery by calculating gridness scores in 30s intervals via a rolling window analysis (Fig. 3d; Supplemental Methods). When normalized against baseline gridness scores, grid cell periodicity was disrupted as early as 9-39 seconds following laser onset (n = 9, normalized gridness score 0.4376 ± 0.2086, One sample Wilcoxon Test; *P*=0.0273), reached its lowest point at 39-69 seconds following laser onset -0.009709 ± 0.3310, *P*=0.0195), and recovered at 60-90 seconds following laser onset (0.62 ± 0.18, *P*=0.0977). These data reveal that disruption of MS-GABAergic input and the reduction of ongoing MEC theta oscillations results in the rapid degradation of grid cell periodicity over a similar time course to that observed in the absence of visual input^4^ and that the grid cell network requires up to 30 seconds to regain stable spatial firing.

**Figure 3.**
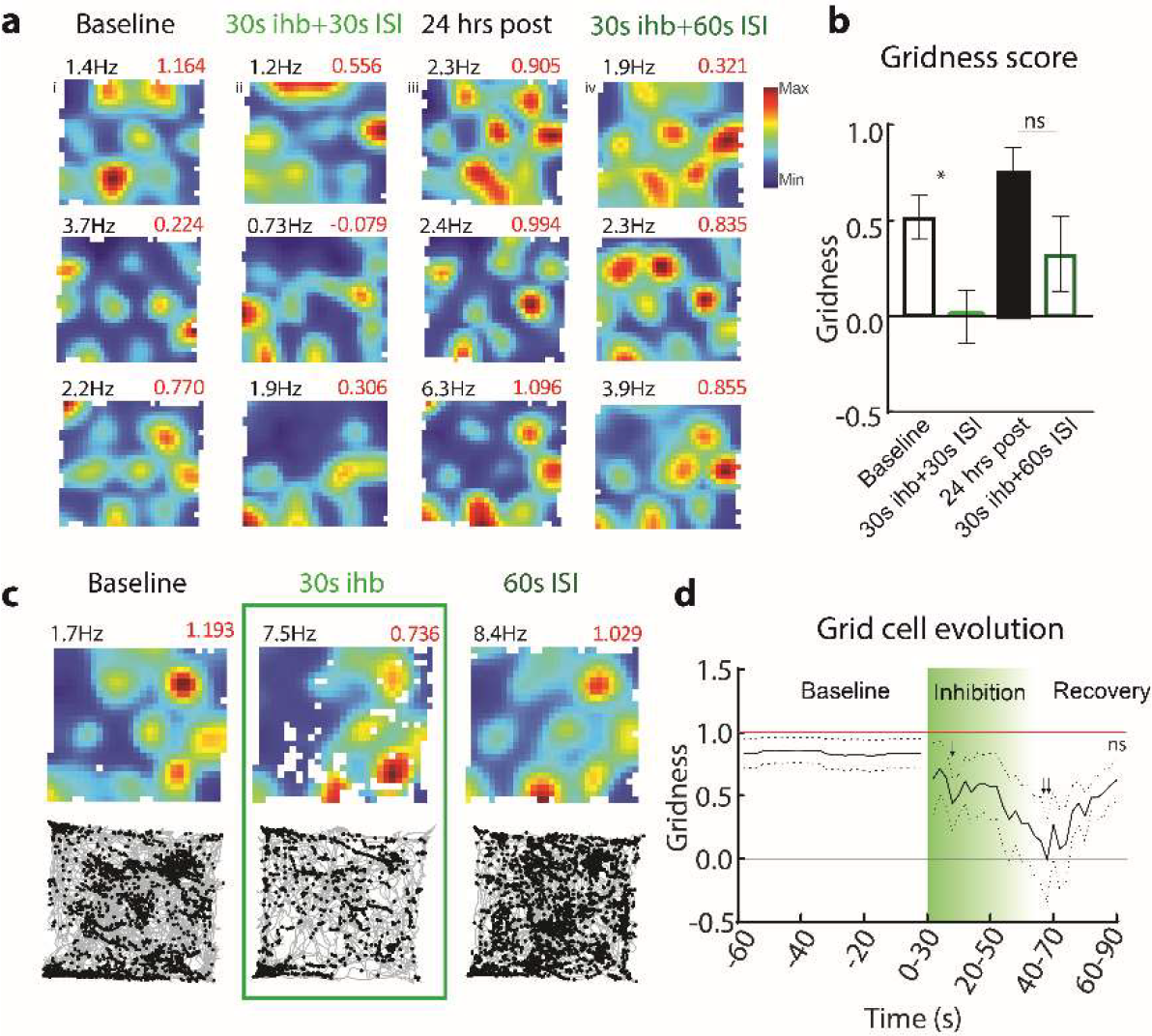
Increasing interstimulus intervals resulted in grid cell spatial periodicity recovery. (a) Example rate maps for cells recorded across i -baseline, ii - combined 30s silencing followed by 30s interstimulus interval (ISI), iii - ∼24 hrs post, iv - 30s silencing periods followed by 60s interstimulus interval stimulation. (b) Grid score for baseline, combined 30s silencing and 30s ISI, Recovery (∼24 hrs), and combined 30s silencing and 60s ISI. (c) →Example rate map and trajectory of baseline and separated rate maps for 30s silencing period and 60s ISI periods. (d) Gridness evolution calculated by a 30 second rolling window beginning one second following stimulation onset normalized to shuffled baseline. For all panels ns = non- significant, \**P*<0.05, → *P*<0.05, →→ *P*<0.01.

Our study supports the hypothesis that grid cell spatial firing relies on theta oscillations and demonstrates the necessity of MS-GABAergic neurons in grid cell spatial and temporal coding. Furthermore, these data reveal that short 30s periods between theta disruptions do not allow grid cell recovery, while a longer 60s window between theta disruption permits the return of grid cell spatial periodicity. Importantly, these data show that while optogenetic inhibition results in an instantaneous disruption to ongoing theta oscillations, it does not immediately degrade grid cell periodicity, but rather the gridness takes up to ∼10-40 seconds to devolve. This mechanism also provides important information on the duration of time that grid cells require to recover following theta disruption (∼60-90 seconds). Our findings provide support to grid cell models where a disruption in representation would take time to recover to a stable attractor space. Together, these data suggest a potential mechanism underlying the spatial navigation and memory deficits related to MS dysfunction.

## Acknowledgements

We would like to thank P. Mannarin and S. Kim for their technical assistance with this project. We are also grateful to J. Q. Lee and M. Yaghoubi who provided comments on prior versions of the manuscript and to all members of the Hasselmo and Brandon laboratories for helpful discussions and advice on this project.

## Data and materials availability

All data, code, and materials information will be made freely available upon publication

## References

1 Hafting, T., Fyhn, M., Molden, S., Moser, M. B. & Moser, E. I. Microstructure of a spatial map in the entorhinal cortex. Nature 436, 801–806, doi:10.1038/nature03721 (2005).

2 Fyhn, M., Molden, S., Witter, M. P., Moser, E. I. & Moser, M. B. Spatial representation in the entorhinal cortex. Science 305, 1258–1264, doi:10.1126/science.1099901 (2004).

3 Moser, E. I., Kropff, E. & Moser, M. B. Place cells, grid cells, and the brain’s spatial representation system. Annu Rev Neurosci 31, 69–89, doi:10.1146/annurev.neuro.31.061307.090723 (2008).

4 Perez-Escobar, J. A., Kornienko, O., Latuske, P., Kohler, L. & Allen, K. Visual landmarks sharpen grid cell metric and confer context specificity to neurons of the medial entorhinal cortex. Elife 5, doi:10.7554/eLife.16937 (2016).

5 Sargolini, F. et al. Conjunctive representation of position, direction, and velocity in entorhinal cortex. Science 312, 758–762, doi:10.1126/science.1125572 (2006).

6 Campbell, M. G. & Giocomo, L. M. Self-motion processing in visual and entorhinal cortices: inputs, integration, and implications for position coding. J Neurophysiol 120, 2091–2106, doi:10.1152/jn.00686.2017 (2018).

7 Winter, S. S., Clark, B. J. & Taube, J. S. Spatial navigation. Disruption of the head direction cell network impairs the parahippocampal grid cell signal. Science 347, 870–874, doi:10.1126/science.1259591 (2015).

8 Bonnevie, T. et al. Grid cells require excitatory drive from the hippocampus. Nat Neurosci 16, 309–317, doi:10.1038/nn.3311 (2013).

9 Brandon, M. P. et al. Reduction of theta rhythm dissociates grid cell spatial periodicity from directional tuning. Science 332, 595–599, doi:10.1126/science.1201652 (2011).

10 Koenig, J., Linder, A. N., Leutgeb, J. K. & Leutgeb, S. The spatial periodicity of grid cells is not sustained during reduced theta oscillations. Science 332, 592–595, doi:10.1126/science.1201685 (2011).

11 Jeffery, K. J., Donnett, J. G. & O’Keefe, J. Medial septal control of theta-correlated unit firing in the entorhinal cortex of awake rats. Neuroreport 6, 2166–2170, doi:10.1097/00001756-199511000-00017 (1995).

12 Mitchell, S. J., Rawlins, J. N., Steward, O. & Olton, D. S. Medial septal area lesions disrupt theta rhythm and cholinergic staining in medial entorhinal cortex and produce impaired radial arm maze behavior in rats. J Neurosci 2, 292–302, doi:10.1523/JNEUROSCI.02-03-00292.1982 (1982).

13 Mizumori, S. J., Perez, G. M., Alvarado, M. C., Barnes, C. A. & McNaughton, B. L. Reversible inactivation of the medial septum differentially affects two forms of learning in rats. Brain Res 528, 12–20, doi:10.1016/0006-8993(90)90188-h (1990).

14 Winson, J. Loss of hippocampal theta rhythm results in spatial memory deficit in the rat. Science 201, 160–163, doi:10.1126/science.663646 (1978).

15 Chrobak, J. J., Stackman, R. W. & Walsh, T. J. Intraseptal administration of muscimol produces dose-dependent memory impairments in the rat. Behav Neural Biol 52, 357–369, doi:10.1016/s0163-1047(89)90472-x (1989).

16 Colom, L. V., Castaneda, M. T., Reyna, T., Hernandez, S. & Garrido-Sanabria, E. Characterization of medial septal glutamatergic neurons and their projection to the hippocampus. Synapse 58, 151–164, doi:10.1002/syn.20184 (2005).

17 Simon, A. P., Poindessous-Jazat, F., Dutar, P., Epelbaum, J. & Bassant, M. H. Firing properties of anatomically identified neurons in the medial septum of anesthetized and unanesthetized restrained rats. J Neurosci 26, 9038–9046, doi:10.1523/JNEUROSCI.1401-06.2006 (2006).

18 Hangya, B., Borhegyi, Z., Szilagyi, N., Freund, T. F. & Varga, V. GABAergic neurons of the medial septum lead the hippocampal network during theta activity. J Neurosci 29, 8094–8102, doi:10.1523/JNEUROSCI.5665-08.2009 (2009).

19 Freund, T. F. & Antal, M. GABA-containing neurons in the septum control inhibitory interneurons in the hippocampus. Nature 336, 170–173, doi:10.1038/336170a0 (1988).

20 Gonzalez-Sulser, A. et al. GABAergic projections from the medial septum selectively inhibit interneurons in the medial entorhinal cortex. J Neurosci 34, 16739–16743, doi:10.1523/JNEUROSCI.1612-14.2014 (2014).

21 Amilhon, B. et al. Parvalbumin Interneurons of Hippocampus Tune Population Activity at Theta Frequency. Neuron 86, 1277–1289, doi:10.1016/j.neuron.2015.05.027 (2015).

22 Dannenberg, H. et al. Synergy of direct and indirect cholinergic septo-hippocampal pathways coordinates firing in hippocampal networks. J Neurosci 35, 8394–8410, doi:10.1523/JNEUROSCI.4460-14.2015 (2015).

23 Zutshi, I. et al. Hippocampal Neural Circuits Respond to Optogenetic Pacing of Theta Frequencies by Generating Accelerated Oscillation Frequencies. Curr Biol 28, 1179–1188 e1173, doi:10.1016/j.cub.2018.02.061 (2018).

24 Lepperod, M. E. et al. Optogenetic pacing of medial septum parvalbumin-positive cells disrupts temporal but not spatial firing in grid cells. Sci Adv 7, doi:10.1126/sciadv.abd5684 (2021).

25 Boyce, R., Glasgow, S. D., Williams, S. & Adamantidis, A. Causal evidence for the role of REM sleep theta rhythm in contextual memory consolidation. Science 352, 812–816, doi:10.1126/science.aad5252 (2016).

26 Dannenberg, H., Kelley, C., Hoyland, A., Monaghan, C. K. & Hasselmo, M. E. The Firing Rate Speed Code of Entorhinal Speed Cells Differs across Behaviorally Relevant Time Scales and Does Not Depend on Medial Septum Inputs. J Neurosci 39, 3434–3453, doi:10.1523/JNEUROSCI.1450-18.2019 (2019).

27 Quirk, C. R. et al. Precisely timed theta oscillations are selectively required during the encoding phase of memory. Nat Neurosci 24, 1614–1627, doi:10.1038/s41593-021-00919-0 (2021).

28 Han, X. et al. A high-light sensitivity optical neural silencer: development and application to optogenetic control of non-human primate cortex. Front Syst Neurosci 5, 18, doi:10.3389/fnsys.2011.00018 (2011).

29 Yartsev, M. M., Witter, M. P. & Ulanovsky, N. Grid cells without theta oscillations in the entorhinal cortex of bats. Nature 479, 103–107, doi:10.1038/nature10583 (2011).

30 Burgess, N., Barry, C. & O’Keefe, J. An oscillatory interference model of grid cell firing. Hippocampus 17, 801–812, doi:10.1002/hipo.20327 (2007).

31 Burgess, N. Grid cells and theta as oscillatory interference: theory and predictions. Hippocampus 18, 1157–1174, doi:10.1002/hipo.20518 (2008).

32 Navratilova, Z., Giocomo, L. M., Fellous, J. M., Hasselmo, M. E. & McNaughton, B. L. Phase precession and variable spatial scaling in a periodic attractor map model of medial entorhinal grid cells with realistic after-spike dynamics. Hippocampus 22, 772–789, doi:10.1002/hipo.20939 (2012).

