## Supplemental Material for "Optogenetic Silencing of Medial Septal GABAergic Neurons Disrupts Grid Cell Spatial and Temporal Coding in the Medial Entorhinal Cortex"

This file includes:

Materials and Methods

Figs. S1-S4

#### Material and Methods

***Subjects.*** All procedures were performed according to protocols and guidelines approved by the Institutional Animal Care and Use Committee at Boston University and the McGill University Animal Care Committee and the Canadian Council on Animal Care. VGAT$^{Cre}$ knock-in homozygote mice (The Jackson Laboratory, stock # 016962) were housed in a 12:12 hour light/dark cycle with food and water ad libitum. The housing room conditions of the mice were maintained at 20–22 degrees Celsius and 21–30% humidity. Specificity of this mouse line for Cre recombinase has previously been confirmed in this mouseline (Boyce et al., 2016).

***Surgeries.*** For all surgeries, mice were anesthetized via oxygen and 5% isoflurane in an inhalation box and then transferred to the stereotaxic frame (David Kopf Instruments), where anesthesia was maintained via inhalation of oxygen and 1-2.5% isoflurane for the duration of the surgery. The animals' body temperature was maintained via heating pad and eyes were protected with hydrogel (Optixcare). The anesthetic Carprofen was administered subcutaneously during each surgery.

To drive the expression of the opsins, VGAT-Cre mice were stereotaxically injected in the MS with a Cre-dependent adeno-associated viral vectors. Silencing experiments were performed with ArchT-GFP (AAVdj-EF1α-Flex-ArchT-GFP from University of North Carolina Virus Core) and virus control experiments were performed with GFP (AAV2-syn-GFP from). Viruses were delivered directly into the medial septum at AP 0.86 mm from bregma, ML 0.0 mm, DV 4.5-4.7mm. All injections were administered via glass pipettes connected to a Nanoject II (Drummond Scientific) injector at a flow rate of 23 nl/sec.

Following a 2-4 week incubation period, mice underwent a second surgery to implant an optic fibre and microdrive. For each mouse, two stainless anchor screws (B000FN0J58, Antrin Online) were placed in front of the inferior cerebral vein, and were secured to the skull with dental cement (Patterson Dental, Inc). A ground wire was positioned above the contralateral cerebellum. For optogenetic modulation of the medial septum an optic fiber (CF230, Thorlabs) connected to a ferrule (Precision Fiber Products or Thorlabs) was implanted at a 5° angle to target just above the medial septum (AP 0.86, ML 0.2, DV 3.83 mm) and secured to the skull with dental cement. Electrophysiological experiments were performed with custom made microdrives. The microdrive used consisted of four independently movable tetrodes (Axona, Inc) and was implanted above of the MEC at the following stereotaxic coordinates: 3.4 mm lateral to the midline, 0.25-0.40 mm anterior to the transverse sinus at an 8-10°angle. Tetrodes were gold-plated using the NanoZ (Neuralynx) to lower impedances to 200-250 kΩ at 1 kHz and dipped in mineral oil prior to surgery. The microdrive was secured in place with metabond and dental cement (Patterson Dental). Once the cement was dry, tetrodes were lowered 400um below the surface of the brain. Mice body temperature was maintained using a heating pad throughout the surgery until fully recovered.

***Data acquisition.*** Following surgery, animals had 1 week of recovery. Once recovered, mice were placed on water restriction and maintained at 85% of their *ad libidum* weight for the duration of experiments. Animals were connected to custom-built headstage preamplifier tethers (Neuralynx) and an optic fiber patch cord (Thorlabs). Light delivery was achieved through an optic fiber patch cord and sleeve coupled to a 530 nm laser (LaserGlow), and light intensity was set for 20 mW. Neural recordings were amplified and band-pass filtered between 0.6 kHz and 6 kHz using a Digital Lynx SX recording system (Neuralynx) and stored with Cheetah Software (Neuralynx). Animals were recorded in an open field (75 x 75 cm square box). Tetrodes were advanced at 25 μm increments to sample neurons. Spike waveform thresholds were adjusted before commencing each recording and ranged between 25-140 μV depending on unit activity. Waveforms that crossed threshold were digitized at 32 kHz and recorded across all four channels of the given tetrode. Local field potentials were recorded across all tetrodes.

**Optogenetic Experiments**. During recording, the environment was dimly lit and white-noise was played to mask uncontrolled ambient sounds. Following each recording the environment was cleaned with disinfectant (Prevail). Once grid cells had been identified during a baseline recording, light delivery was achieved through an optic fiber patch cord and sleeve coupled to a green laser (520nm wavelength, maximum power output: 60mW, Doric Lenses) and light intensity at the tip of the optic fiber was estimated for between 15-20 mW. Stimulation protocols consisted of 30s laser-on periods followed by 30s laser-off periods.

**Histology.** Immunocytochemistry was used to assess the virus distribution across the septum and localize tetrode recording sites in the MEC. Experimental animals were euthanized following experiments. Mice were anesthetized and intracardially perfused with 4% paraformaldehyde in PBS. Brains post-fixed by immersion in the same fixative for 2 weeks with the tetrodes in place following perfusion and then dissected from the skull. Free-floating coronal sections of the entire medial septum and sagittal sections of the entire entorhinal cortex were cut using a vibratome (40 µm) or on the cryostat (25 µm). Free-floating sections across the septum were incubated in GFP rabbit antiserum (1:1000, Invitrogen), and were detected with anti-rabbit coupled to AlexaFluor-488 (1:1000, Invitrogen). Sections of the entorhinal cortex were stained for DAPI to located tetrode tips. The slices were mounted with Fluoromount-G (Southern Biotechnology) and analyzed with an AxioObserver.Z1 microscope (Carl Zeiss).

***Data processing.***

**Cluster cutting.** Single-units were isolated 'offline' manually using graphical cluster cutting software (Offline Sorter, Plexon Inc.) individually for each recording session. Neurons were separated based on the peak amplitude and principal component measures of spike waveforms. Stability of units was confirmed by tracking waveform profiles across the four leads of the tetrode and cluster position across recording sessions, comparing to the baseline session.

**Position estimation.** To estimate the position of the animal, we measured the centroid of a group of red and green diodes positioned on the recording head stage. Head direction was calculated as the angle between the red and green diodes.

***Data analysis.*** To analyze LFP power, the power spectrum for local field potentials was obtained using multitaper method included in the Chronux toolbox (mtspectrumc with NW = 3 and K = 5; (Mitra & Bokil, 2007). Theta power was calculated by taking the area within 1Hz of the maximum power in the theta range (6-10Hz). Baseline theta power was obtained by taking the average of theta power during laser-off trials, and mean reduction in theta power for each session was obtained by dividing each laser-on trial with baseline theta power and averaging across laser-on trials. For each channel, the theta-delta ratio was obtained by taking the ratio between mean theta power (6-10Hz) and mean delta power (2-4Hz) during laser-off segments. The channel with the highest theta-delta ratio was used in the LFP analysis.

**Measurement of single unit properties.** Spatial rate maps were constructed for a given unit by taking the number of spikes in 3.6 cm by 3.6 cm spatial bins and dividing by the amount of time spent in that bin. Rate maps were smoothed using a 5x5 bin 2D-Gaussian kernel with a one-bin standard deviation. The spatial periodicity of grid cells was quantified with a ‘‘gridness” score and computed from the spatial autocorrelation of the smoothed rate maps (Brandon et al., 2011). In brief, to calculate the gridness score, ‘‘gridness 3” from Brandon et al. (2011), the center peak was removed from the autocorrelation and if present, the six surrounding peaks were found and cut out to make a donut. If the grid shape was elliptical, it was distorted to create a circle. The correlation was calculated for each 3-degree rotation of the donut to itself. The gridness score is the difference between the correlation at the minimum peak of 60 or 120 degrees and the maximum trough at 30, 90, or 150 degrees. In line with Dannenberg et al., 2020, grid cells were defined as cells with a grid score >=0.19 and at least 300 spikes in the baseline condition. The spatial information of a cell was calculated using the same rate map as above, with the equation for *I* in bits/spike where $p_i$ is the probability of occupancy in pixel *i,* $F_i$ is the firing rate for pixel *I,* and *F* is the mean firing rate.

Gridness evolution was calculated using a 30-second rolling window with 1-second steps beginning at 1 second following later onset. Baseline grid score values were calculated using 50 shuffled start times and both the recovery and baseline values were averaged over 2 second periods.

***Statistics***. All statistical tests are noted where the corresponding results are reported throughout the main text and supplement. All tests were 2-tailed unless otherwise noted. Nonparametric tests were used as neural data violates the assumptions of parametric tests.

Statistical analysis: All statistical evaluations were performed under MATLAB and Prism. Non-parametric Wilcoxon signed rank tests, Wilcoxon rank sum tests, Kolmogorov-Smirnov tests, and Kruskal-Wallis or Friedman test one-way ANOVAs were used throughout the paper. P-values reported from all post hoc tests were with Dunn's multiple comparison test.

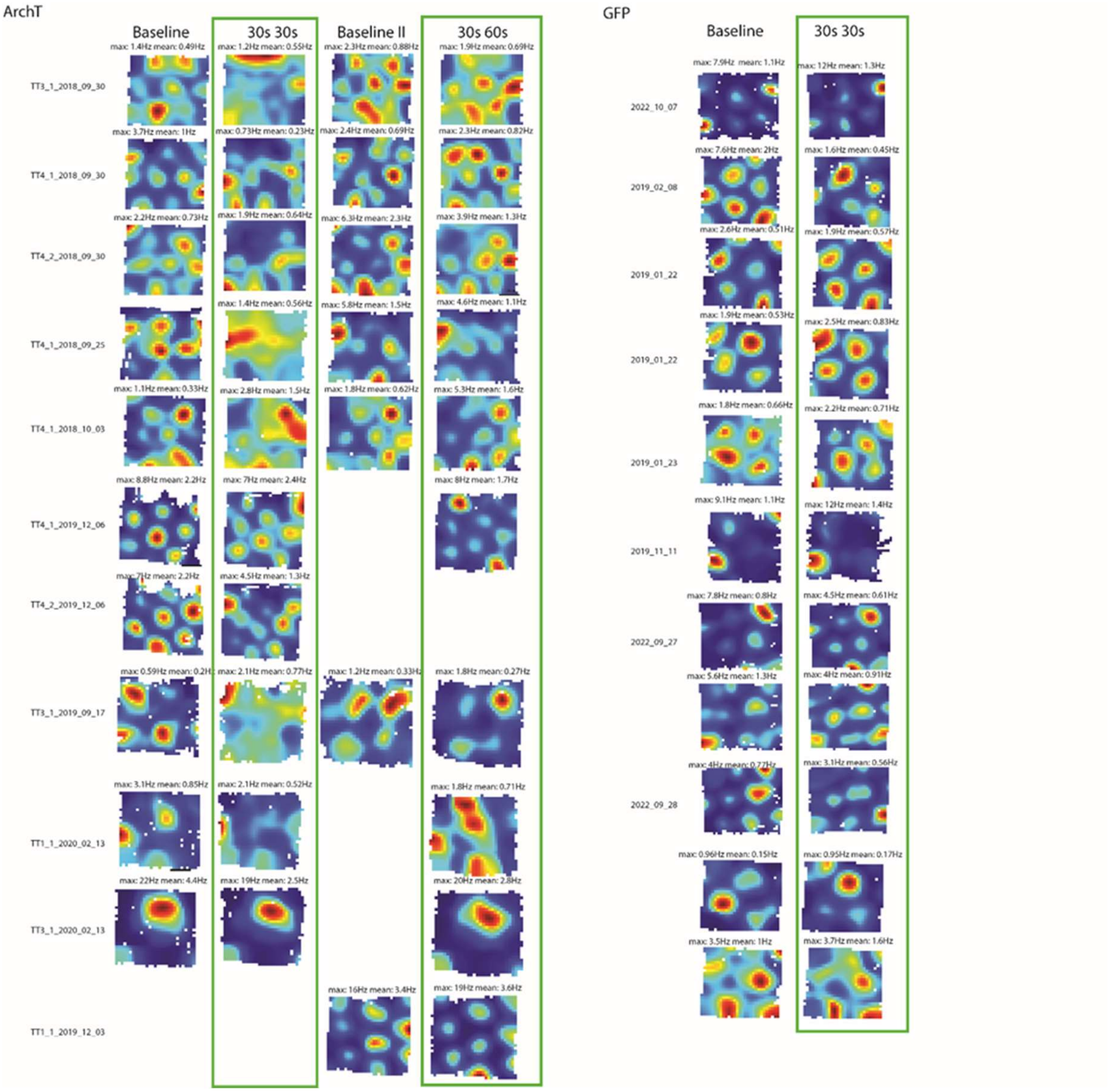


**Fig. S1**. Overview of grid cells recorded during baseline, 1st stimulation of 30s inhibition (ihb)-30s interstimulus interval (ISI), recovery and 2nd stimulation sessions of 30s ihb-60s ISI. Rate maps are provided for each unit. Max and mean firing rate is shown above each rate map. Note that the color-coding of the rate maps are relative to each recording session for each unit. Tetrode and recording date is indicated to the left of the rate maps. Experimental paradigm is shown above each column and optogenetic stimulation sessions are marked by green boxes.

### GFP Control

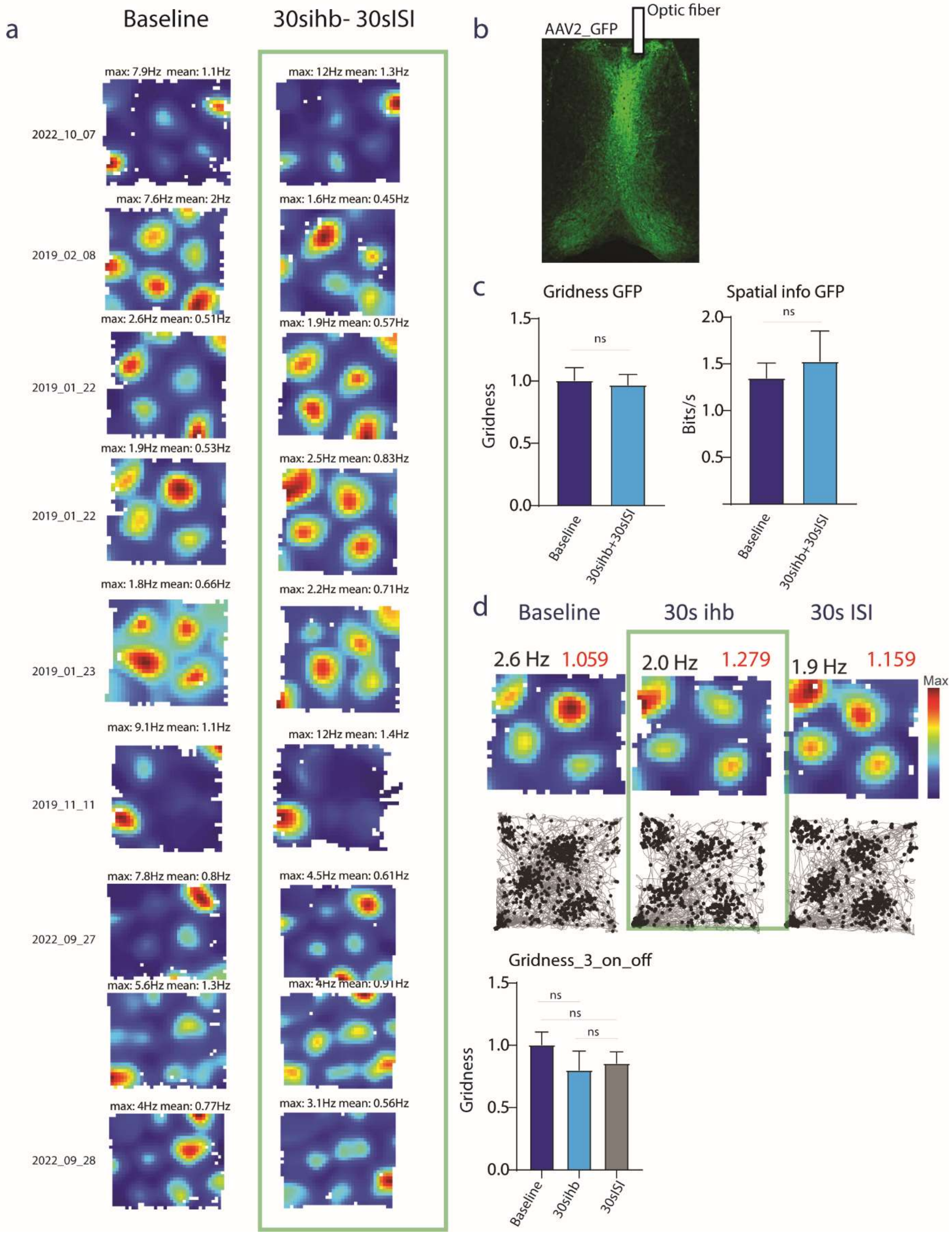

a
Baseline
30sihb- 30sISI
max: 7.9Hz mean: 1.1Hz
max: 12Hz mean: 1.3Hz
2022_10_07
max: 7.6Hz mean: 2Hz
max: 1.6Hz mean: 0.45Hz
2019_02_08
max: 2.6Hz mean: 0.51Hz
max: 1.9Hz mean: 0.57Hz
2019_01_22
max: 1.9Hz mean: 0.53Hz
max: 2.5Hz mean: 0.83Hz
2019_01_22
max: 1.8Hz mean: 0.66Hz
max: 2.2Hz mean: 0.71Hz
2019_01_23
max: 9.1Hz mean: 1.1Hz
max: 12Hz mean: 1.4Hz
2019_11_11
max: 7.8Hz mean: 0.8Hz
max: 4.5Hz mean: 0.61Hz
2022_09_27
max: 5.6Hz mean: 1.3Hz
max: 4Hz mean: 0.91Hz
max: 4Hz mean: 0.77Hz
max: 3.1Hz mean: 0.56Hz
2022_09_28
b
AAV2_GFP
Optic fiber
c
Gridness GFP
Spatial info GFP
ns
Gridness
Bits/s
0.0
0.5
1.0
1.5
2.0
Baseline
30sihb+30sISI
d
Baseline
30s ihb
30s ISI
2.6 Hz 1.059
2.0 Hz 1.279
1.9 Hz 1.159
Max
Gridness_3_on_off
Baseline
30sihb
30sISI

**Fig. S2**. GFP control. A. Rate maps are shown for multiple different neurons (rows) for the baseline condition (left) and the laser-on period (30s inhibition (ihb)-30s ISI) marked by the green box. Note that the light on will not evoke inhibition in the GFP controls and does not alter grid cell firing. B. Example of GFP virus expression across the MS. C. Comparison of gridness score and spatial information during baseline and stimulation periods. D. Top. Comparison of recordings during baseline, 30s laser-on and 30s ISI periods. Bottom. Gridness score shows no significant change in the GFP controls.

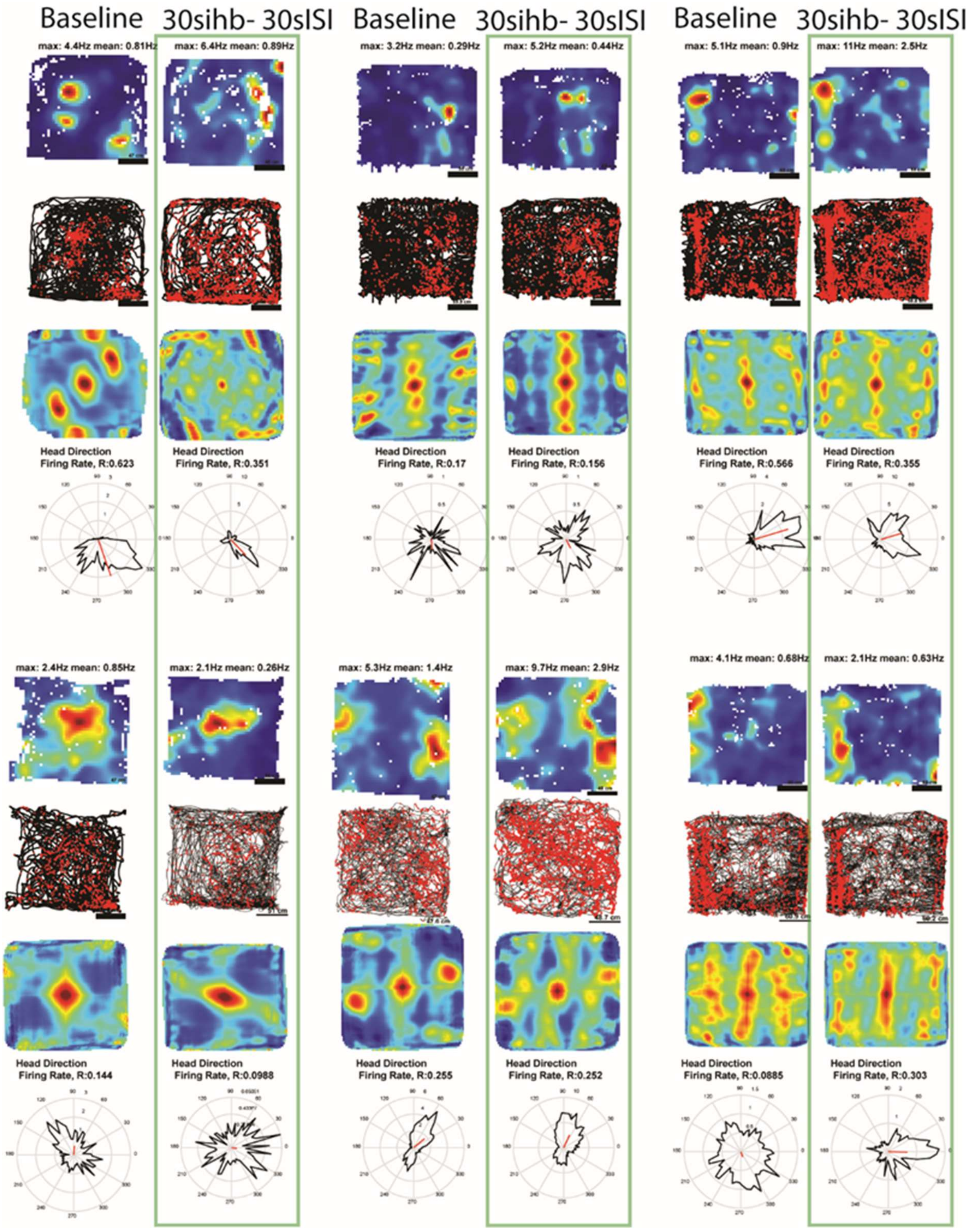


**Fig. S3**. Examples of spatially modulated non-grid cells recorded during baseline and stimulation conditions (these did not pass the gridness-score criterion to be identified as grid cells). Individual rows show: 1) spatial plots of firing rate, 2) animal trajectory with red dots showing location of mouse during each spike 3) spatial autocorrelations of firing rate map, 4) polar plot of firing rate of head directions.

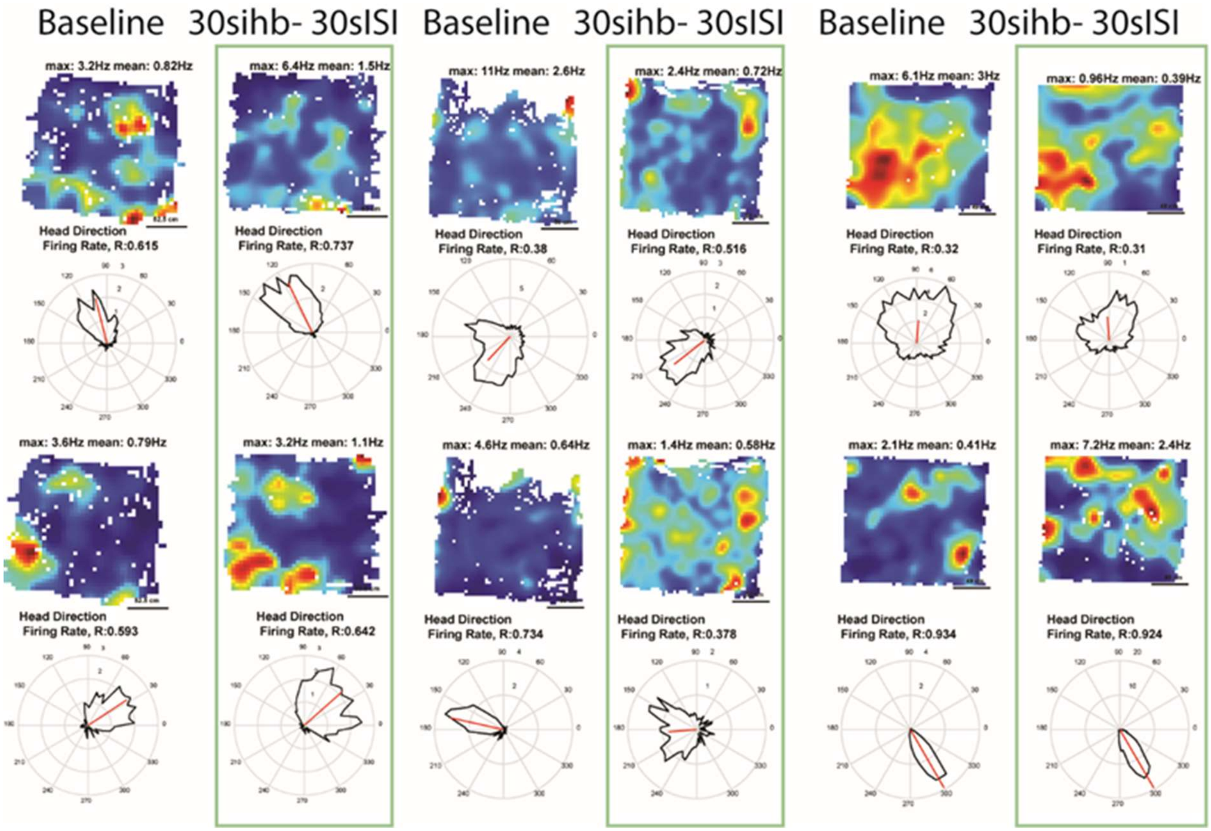


**Fig. S4**. Examples of head direction cells recorded during baseline and 30s ihb-30s ISI conditions. Individual rows show: Top) spatial plots of firing rate, Bottom) polar plot of firing rate over head directions. Note sparing of head direction selectivity during the optogenetic stimulation 30s ihb-30 ISI.
